# Effect of habitat degradation on hantavirus infection among introduced and endemic small mammals of Madagascar

**DOI:** 10.1101/2023.12.24.573235

**Authors:** Jérémy Dubrulle, Kayla Kauffman, Voahangy Soarimalala, Toky Randriamoria, Steven M. Goodman, James Herrera, Charles Nunn, Pablo Tortosa

## Abstract

Hantaviruses are globally distributed zoonotic pathogens capable of causing fatal disease in humans. Rodents and other small mammals are the typical reservoirs of hantaviruses, though the particular host varies regionally. Addressing the risk of hantavirus spillover from animal reservoirs to humans requires identifying the local mammal reservoirs and the predictors of infection in those animals, such as their population density and habitat characteristics. We screened native and non-native small mammals and bats in northeastern Madagascar for hantavirus infection to investigate the influence of habitat, including effects of human land use on viral prevalence. We trapped 227 bats and 1663 small mammals over 5 successive years in and around Marojejy National Park across a range of habitat types including villages, agricultural fields, regrowth areas, and secondary and semi-intact forests. Animals sampled included endemic tenrecs (Tenrecidae), rodents (Nesomyidae) and bats (6 families), along with non-native rodents (Muridae) and shrews (Soricidae). A hantavirus closely related to the previously described Anjozorobe virus infected 9.5% of *Rattus rattus* sampled. We did not detect hantaviruses in any other species. Habitat degradation had a complex impact on hantavirus prevalence in our study system: more intensive land use increase the abundance of *R. rattus*. The average body size of individuals varied between agricultural and non-agricultural land-use types, which in turn affected infection prevalence. Smaller *R.rattus* had lower probability of infection and were captured more commonly in villages and forests. Thus, infection prevalence was highest in agricultural areas. These findings provide new insights to the gradients of hantavirus exposure risk for humans in areas undergoing rapid land use transformations associated with agricultural practices.

## Introduction

Zoonotic diseases represent a major threat to human health, with hundreds of thousands of deaths and millions of infections occurring annually (Cascio et al., 2011). Of the many zoonoses that impact humans, hantaviruses are a growing threat due to their increasing global incidence and high mortality rate (Jonsson et al., 2010). Particularily, the Old World hantaviruses are responsible for more than 100,000 infections per year (Avšič-Županc et al., 2019), and disproportionately affect low-income, rural areas (Jonsson et al., 2010). Of the Old World strains, Seoul *Orthohantavirus* has caused more than 1.6 million infections in China since the 1950’s (Jonsson et al., 2010).

Humans are primarily exposed to hantaviruses through aerosolized excrement of infected rodents, which is broadly associated with activities that bring humans into contact with these animals and where rodent densities are relatively high (Watson et al., 2014). In the Old World hantaviruses, the disease associated with infection in humans is hemorrhagic fever with renal syndrome and has a mortality rate of 15% (Schmaljohn and Hjelle, 1997). Antiviral drugs and vaccines are not yet available, with control instead relying on exposure prevention (Brocato and Hooper, 2019). Thus, understanding the enzootic ecology of these viruses, factors facilitating spillover into non-reservoir hosts, as well as viral evolution, are important aspects to prevent human infections (Jonsson et al., 2010).

*Orthohantavirus* species from the Hantaviridae family form a diverse genus of RNA viruses with a vast set of enzootic reservoir host Old World hantaviruses are primarily harbored by Muridae (rodents) and *Myodes* (voles) (Jonsson et al., 2010). Other reservoirs include shrews (Soricidae) in Guinea (Guo et al., 2013; Klempa et al., 2007), China (Guo et al., 2013), Korea (Lee et al., 2017), Vietnam (Song et al., 2007), and Hungary (Kang et al., 2009b) and bats in China (Guo et al., 2013), Côte d’Ivoire (Sumibcay et al., 2012) and Sierra Leone (Weiss et al., 2012). Phylogenetic studies reveal co-divergence, reassortment, and host-switching between members of the Cricetid rodents, particularily *Lemmus sibirica* and *Microtus fortis* (Vapalahti et al., 1999) and between the Talpidae and Soricidae families (Kang et al., 2009), all of which account for the high diversity and worldwide distribution of hantaviruses.

Climatic factors such as fluctuations in temperature and rainfall in combination with local human modifications in landscape structure affect reservoir abundance [reviewed in (Prist et al., 2017)], which in turn changes the dynamics of hantavirus prevalence and human exposure risk (Klempa, 2009). Abundance of reservoir hosts and infection prevalence varies seasonally with food availability (Nsoesie et al., 2014; Prist et al., 2017). Habitat availability (Suzán et al., 2008) and fragmentation (Fialho et al., 2019; Ganzhorn, 2003; Goodin et al., 2006) also impact community composition, with fragmented areas and agricultural fields often having denser populations of introduced commensal species such as mice (*Mus musculus)* and rats *(Rattus norvegicus and R.rattus.)* (Umetsu and Pardini, 2007). Deforestation also benefits invasive generalist species such as introduced rodents, which in turn tends to decrease the diversity of endemic and native species in the local mammal communities (Pardini et al., 2010).

Madagascar provides a valuable context to investigate the effect of anthropogenic disturbance on hantavirus prevalence in small mammals as the island is experiencing rapid deforestation and habitat fragmentation (Vieilledent et al., 2018). Madagascar’s exceptional biodiversity and levels of endemism are due to isolation in deep geological time (Antonelli et al., 2022). This landmass is home to a unique native small mammal fauna of Madagascar, including 46 species of bats (80% endemic), 31 species of endemic tenrecs (Geogalinae, Oryzorictinae and Tenrecidae), and 28 species of endemic rodents (subfamily Nesomyinae) (Goodman, 2022). Non-native Muridae rodents (*Rattus* and *Mus*) and Soricidae shrews (*Suncus*) also inhabit the island. This isolation and diversity may explain the evolution of unique hantavirus lineages described from the island such as Anjozorobe virus (ANJZV), a variant of Thailand virus (*Orthohantavirus thailandense*), which was found in introduced *Rattus rattus* and endemic *Eliurus majori* on Madagascar (Filippone et al., 2016; Kikuchi et al., 2021; Raharinosy et al., 2018a; Reynes et al., 2014). Human exposure to ANJZV in Madagascar is estimated at 2.7% nationally, with higher seroprevalence (7.2%) in communities adjacent to forests (Rabemananjara et al., 2020). A closely related hantavirus lineage, called *MayoV*, has also been described in *R. rattus* on the neighboring island of Mayotte (Filippone et al., 2016).

Here, we investigated the ecology of hantavirus infection in rural northeastern Madagascar in relation to mammalian community composition and human land use patterns. We surveyed a wide range of terrestrial small mammals and bats for hantavirus over a 5-year period along gradients of land use in and around Marojejy National Park in northeastern Madagascar. We focused on land-use gradients spanning three villages in an area of mixed human activities, including swidden agriculture that results in the clearing of forests and a fragmented landscape. Remnant forests are largely restricted to protected areas, and secondary forests are embedded in a matrix of brushy regrowth and agricultural fields (e.g. rice fields, vanilla agroforests). Previous work showed that these habitat modifications have resulted in higher abundances of commensal small mammals, specifically non-native rodents and shrews around one of our study villages, as compared to the nearby protected forest (Herrera et al., 2020). These factors are anticipated to influence hantavirus ecology by modifying abundance and diversity of mammal hosts and by increasing the risk of contact with hantavirus-infected substrates and materials [reviewed in (Tian and Stenseth, 2019)].

As found in other tropical settings, we hypothesized that hantavirus infections would be more common during the rainy season, driven by increased population densities of small mammal hosts at this time of year (Scobie et al., 2023). We also hypothesized that viral prevalence in small mammals would be higher in large, adult males, as observed in other studies, which attribute this observation to increased direct competition for resources (Jonsson et al., 2010). We expected that viral prevalence would increase with anthropogenic disturbance and agricultural use, as a result of increased *R.rattus* abundance. Lastly, we used the variability in hantavirus prevalence across a range of lande use types to better understand the findings of Raharinosy et al. (2018). reporting lower viral prevalence within houses than outside of them, on Madagascar.

## Material and methods

### Ethics statement and sample collection

Lung tissue samples from wild small mammals were collected between 2017 and 2021 in the vicinity of three villages adjacent to Marojejy National Park in the SAVA Region in northeastern Madagascar, following a standard grid-trapping protocol detailed below. The village of Mandena (14.477049° S, 49.8147° E) and its surroundings were sampled during the dry season (September-December) in 2017 and 2019, during the wet season (March-May) in 2020, and in the transitional dry-wet season (June-August) in 2018 and 2020. A second village, Manantenina (14.497213° S, 49.821347° E), was sampled only in the transitional season (2019) while a third village, Sarahandrano (14.607567° S, 49.647759° E), was sampled during the dry (2020) and wet (2021) seasons. Bats were trapped along flyways located on or adjacent to the trap grids and similar habitat types using harp traps and mist nets. Bats were also captured from caves in the area using butterfly nets. Bat trapping occurred during September-November 2019 in Mandena (dry season) and March-May 2021 in Sarahandrano (wet season).

Small mammal traps were installed in different land-use types, including semi-intact forest within the national park, as well as secondary forest and agroforest outside the park, and agricultural fields, flooded rice fields, and brushy regrowth (fallow areas after swidden cultivation) around villages. Traps were also placed in houses within villages adjacent to the trap grids. Sampling methods varied by location and trap grid; trap grids prior to September 2019 were 90 m^2^ and after that date were 100 m^2^ and consisted of metal (Sherman, model LFAm Tallahassee, Florida, USA) and mesh (Tomahawk, model 201, Hazelhurst, Wisconsin, USA) live traps placed 10 m apart. At each sampling site, two pitfall lines of 100 m in length, composed of 15 L buckets placed 10 m apart and the line bisected by a vertical plastic drift fence partially buried. The pitfall lines were located 20 to 50 m from the trap grid. Most grids were sampled for six consecutive nights, but where abundance was low, longer trapping periods were needed (up to 15 nights), and trapping within houses was limited to five nights. All non-native animals and a subset of native animals were collected. Tissue samples were stored in 70% ethanol and placed in long term storage for 1-4 months at -20°C until the molecular analyses were conducted.

All animals were processed using the same methods and samples were stored in the same conditions until further laboratory analysis. All procedures were approved by IACUC at Duke University (protocol number A002-17-01, 2017–2019, A262-19-12 2019-2021) and by Malagasy authorities (No. 289/17, 146/18, 280/19, 57/20, 191/20, 307/21— MEEF/SG/DGF/DSAP/SCB).

### Nucleic acids extraction

Lung samples were rehydrated overnight at 4°C in 1.5 mL of autoclaved milliQ water. Then 25-50 mg of rehydrated tissue was cut into small pieces with a sterile disposable scalpel blade and transferred into individual 2 ml Eppendorf tubes containing 180 µL of ATL and 20 µL of proteinase K provided by the IndiSpin® QIAcube® HT Pathogen Kit (Qiagen, Courtaboeuf, France). Tubes were incubated between 6 and 12 h at 56°C until full proteolysis was achieved. Extraction was then performed in 96 well plates using the Qiagen Cador pathogen kit and QiaCube XT robot following the manufacturer’s instructions. Nucleic acids were collected in a 200 µL elution buffer and stored at -80°C.

### Hantavirus detection

Reverse transcription was conducted with 10 µL of eluted nucleic acids using the ProtoScript® II Reverse Transcriptase (New England Biolabs, Massachusetts, USA). For each sample, RNA was denatured at 70°C for 5 min. in a mix containing 1 µL of DNTP, 1 µL of Rnase free water and 0.5 µL of random primers and thawed in a cold block. Then, a second mix was added, containing 4 µL of protoscript buffer, 2 µL of 10X DTT, 1 µL of the reverse transcription enzyme, and 0.5 µL of RNAsin (New England Biolabs, Massachusetts, USA), resulting in a total volume of 20 µL. The mix was incubated at 25°C for 10 min., 42°C for 50 min. and 65°C for 20 min.

cDNAs were used as a template in a previously published nested-PCR targeting the L-segment coding for the RNA-polymerase RNA-dependent (Klempa et al., 2006). The mix contained 12.5 µL of GoTaq® G2 Hot Start Polymerase (Promega,, Wisconsin, USA), 1 µL of degenerated primers at 10 µM (HAN-L-F1 and HAN-L-R1 for primary PCR, HAN-L-F2 and HAN-L-R2 for secondary PCR). 2 µL of cDNA was used for the primary PCR’s template. The secondary PCR was prepared using 0.5 µL of the primary PCR product as DNA template. Thermal cycling conditions were identical for both primary and secondary PCR and were as follows: nucleic acids were denatured at 95°C for 5 min. followed by 2 cycles at 94°C for 45 sec., 46°C for 45 sec. and 72°C for 60 sec., 2 cycles at 94°C for 45 sec., 44°C for 45 sec. and 72°C for 60 sec., and then 30 cycles at 94°C for 45 sec., 42°C for 45 sec. and 72°C for 60 sec. before finalizing the PCR with 72°C for 10 min. DNA was visualized using a 1.8% TBE agarose gel stained with Gel Red (Biotium, Fremont, California, USA).

### PCR targeting hantavirus in endemic hosts

Additional primers were developed to detect hantavirus RNA that might be hosted by endemic Malagasy small mammals. Primers were either referenced from the literature (Table S1) or newly designed using available reference sequences of ThaiV (MZ343362.1), MayoV (KU587796.1) and ANJZV (LC553724.1, NC_034556.1, KC490924.1, KC490923.1 and KC490922.1). Three semi-nested PCR schemes were used to screen endemic mammals and their associated nucleotide sequences are presented in Table S1. The lack of positive hantavirus detection in endemic mammals resulted in the need to test newly designed primers on positive *R. rattus* found in this study, through a nested PCR protocol. Primer pairs that did not successfully amplify these *R. rattus* hantaviruses were excluded from subsequent analyses, resulting in three semi-nested PCR systems, which were all used following these PCR conditions: initial denaturation at 95°C for 5 min. was followed by 35 cycles of 45 sec. at 95°C, 45 sec. annealing at 50°C and 1 min. elongation at 72°C. PCR ended with a final elongation step of 10 min. at 72°C. PCR products were visualized on a 2% agarose gel and gel bands with the expected size were gel purified (QIAquick Gel Extraction Kit) and used for Sanger sequencing (Genoscreen, Lille, France).

### Phylogenetic analyses

Positive nested PCR samples were Sanger sequenced on both DNA strands at Genoscreen (Lille, France). Chromatograms were manually edited using Geneious 9. Sequences of the partial L segment, 347 bp, are available on GenBank under accession numbers OP328829-OP328903 (Table S5). We included in the analyses sequences from reference HantaV strains belonging to lineages directly related to ThaiV viruses, and using Sangassou virus as an outgroup. The final tree was based on a set of 85 sequences of 347 bp composed of 11 external reference sequences related to ThaiV hantavirus, and 74 sequences generated in the context of this study. We imported our dataset in Datamonkey Adaptive Evolution server to identify the presence of recombinant strains (https://www.datamonkey.org/). The most appropriate phylogenetic model was determined using PhyML3. Phylogenetic signal was checked for with DAMBEE and the tree was built on 20,000,000 iterations. MEGA10 was used to select the appropriate evolution model. MrBayes package on Geneious 9.1 was fed with all aforementioned settings while other settings were set to default values. Reference sequences were trimmed to 347pb and all sequences were manually aligned using the “Maft alignment” plugin on Geneious 9.

### Statistical Analyses

All analyses were performed in R version 4.3.0 (R Core Team, 2023). All pairwise comparisons were made using a significant level of 0.05. Only *R. rattus* were considered in the analyses as hantaviruses were only detected in this species. We used generalized linear mixed-effects models (GLMMs) with a binomial error structure to investigate the effect of environmental and individual variables on hantavirus infection. To examine the random effect of trap grid identity, a unique identifier given to each trap grid installation was included in all models to control for the non-independance of animals captured in the same grid. We considered the fixed environmental effects of season (dry, wet, and transitional), village, direct trap distance to the village, and habitat type (semi-intact forest, secondary forest, agroforest, brushy regrowth, agriculture, flooded rice fields and village). Individual-level fixed effects considered were sex, age based on tooth eruption and wear (sub-adult and adult), mass (g), head-body length (mm), body condition score, and the interaction term between mass and head-body length. The body condition was calculated using mass per length cubed (Fulton’s index) and served as an indicator for an individual’s health. A final fixed effect, the number of *R. rattus* captured during each trap grid installation (e.g. flooded rice field in Mandena during the dry season 2019), was also used as an approximation for animal density. To assess whether variation in hantavirus prevalence between habitat types could be explained by individual-level traits, environmental traits, or rat density, we used a model containing the combinations of those traits (environmental + demographic, environmental + number of *R. rattus*, and environmental + demographic + number of *R. rattus*). The data used in all models was subset to include observations containing all representative metadata. Furthermore, trapping effort and sampling methods in villages differed substantially from all other sites. Thus, these variables were excluded in the models that included animal density. All models were also rerun with the complete data set available for covariates included to verify the robustness of the results.

Global models were fitted using the glmmTMB package, version 1.1.7 (Brooks et al., 2017) and diagnostic tests were performed using the DHARMa package, version 0.4.6 ((Hartig, 2020). We used the MuMIn package, version 1.47.5 (Barton 2023) ‘dredge’ function to identify the best models based on the corrected Akaike information criterion (AICc), then retained all models in the 95% cumulative sum weight confidence set to find the cumulative sum AICc weights (importance) for each predictor and calculate model-averaged parameter estimates. We used the full averaged model to perform Tukey-adjusted post-hoc pairwise comparisons using the emmeans package, version 1.8.6 (Lenth 2023). The proportion of variance explained by the marginal (fixed) effects of best model (pseudo R2) was found using the ‘r.squaredGLMM’ function in the MuMIn package (Barton, 2023).

## Results

### 1.1. Animal composition and hantavirus prevalence

We trapped 1681 small mammals, of which 1663 were tested for hantavirus infection. The tested samples consisted of 72.94% (N=1213) nonnative and 27.1% (N=450) endemic mammals (Table S2). Introduced *R. rattus* represented 47.8% (N=794) of the tested animals and the only species in which hantavirus was detected. We thus limited the statistical analyses to this species.

Hantavirus prevalence in *R. rattus* was 9.5% (75/794, 95% CI: 7.5-11.7%) and varied with environmental and demographic factors as well as animal density (Fig. 1, Table S3). The age composition varied significantly by habitat type (Fisher’s exact test, p<0.001; Fig. 1a). The average body mass of infected rats (124.97 g) was significantly greater than uninfected individuals (81.70 g; Fig. 1b; Welch’s two-sample t-tests, p<0.001). The average mass of *R. rattus* varied significantly by habitat type (Fig.1c; Kruskal-Wallis, p<0.001) and the infected individuals in each habitat type were mostly of above average mass. The same trends displayed in (Fig. 1b) and (Fig. 1c) were observed for head-body length (mm) but not body condition (g/mm3); see Fig. S1 and Fig. S2.

**Figure 1:**
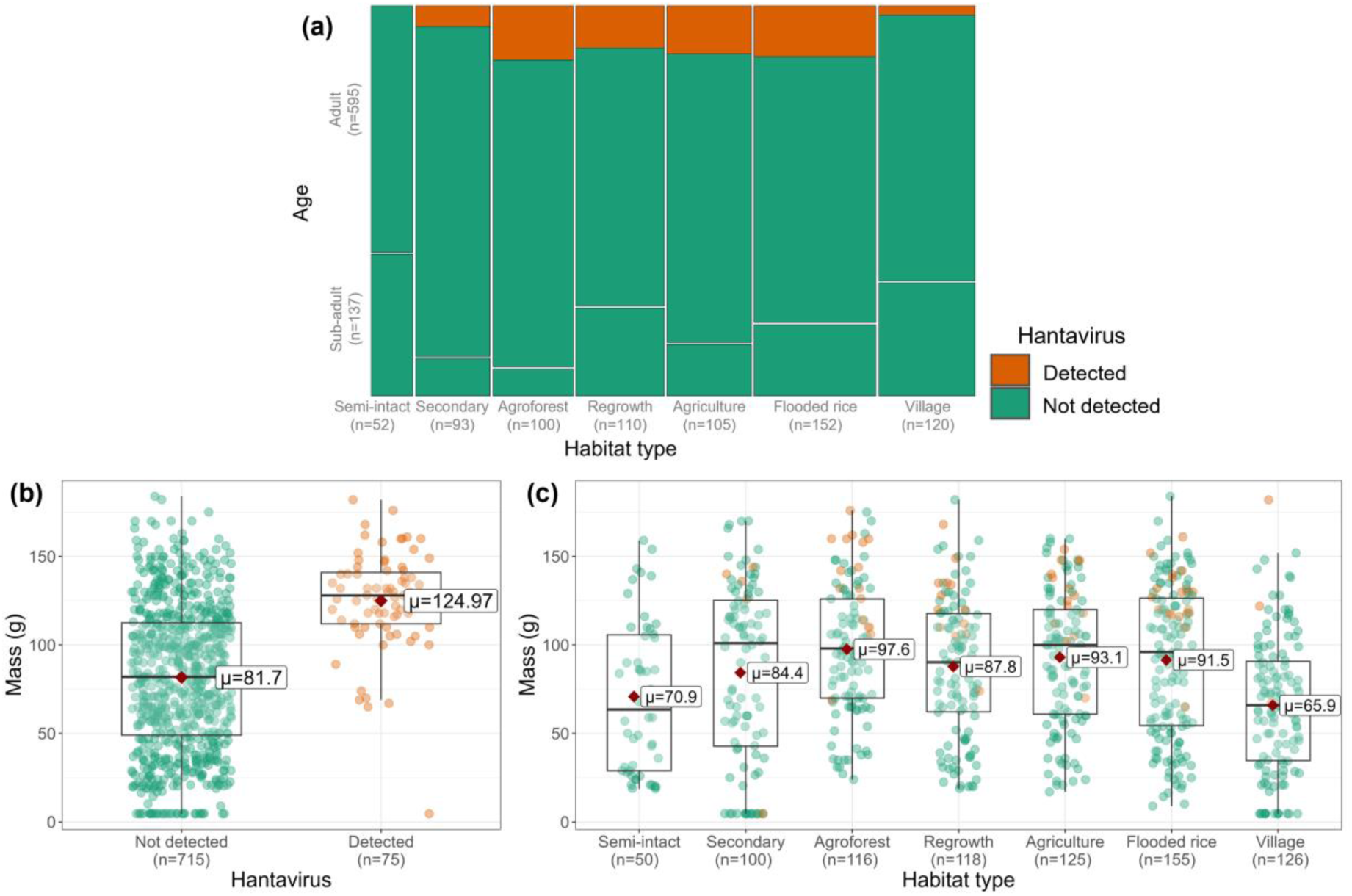
Hantavirus prevalence by (a) age class, (b) body mass, and (c) mass and habitat type. Orange represents animals in which hantavirus was detected and green not detected. The average (µ) mass (g) is displayed on (b) and (c) and indicated with a red diamond. Individuals with missing age information (n=62) were omitted from plot (a) and individuals with missing mass information (n=4) were omitted from the plot (b) and (c).

### 1.2. Complementary screening of endemic mammals

All endemic species of small mammals and bats tested negative through nested PCR (Klempa et al., 2006). To test for possible false negative PCRs resulting from infections with distinct hantaviruses, we tested most endemic mammals, as well as introduced shrews, with alternative PCR schemes. For this, we first validated alternative PCR schemes with known positive *R. rattus* from this study. Seven out of 16 primers produced amplicons of the expected size and were further tested on subsequent endemic rodents. Samples from 214 endemic mammals representing 12.8% of all trapped mammals including *Microgale brevicaudata* (N=176), *Eliurus* spp. (N=9), *Setifer setosus* (N=29), *Suncus murinus* (N=55), and native bats (N=141, see Table S1) were tested. None of the samples tested positive using these alternative end-point PCR schemes.

## 2. Genetic diversity of the newly found hantavirus

All amplicons matching the expected size were sequenced, resulting in partial L segments of 347bp. Genetic identity with previously reported hantaviruses from Madagascar (Kikuchi et al., 2021; Raharinosy et al., 2018a; Reynes et al., 2014b) ranged between 85.0% and 91.6%. Identity restricted to our study cohort only (75 sequences) ranged from 89.6% to 100%. Forty one of the 75 obtained sequences were unique, indicating a fairly high level of diversity. Our sequences did not form a phylogenetic cluster with hantaviruses known from nearby islands (Fig. 2).

**Figure 2.**
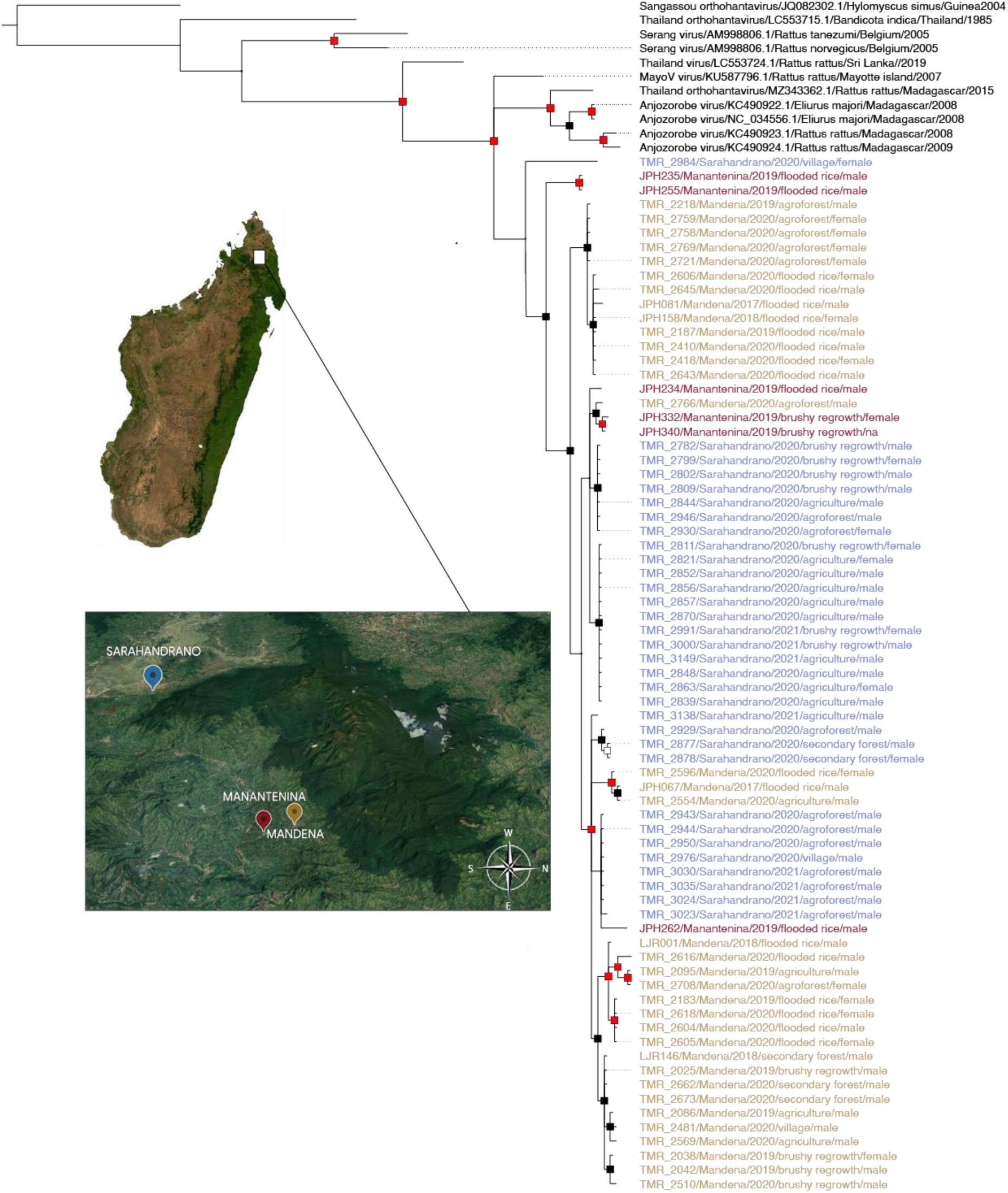
Bayesian phylogeny of sequence hantavirus samples. Bayesian phylogeny showing relatedness of hantaviruses from within and near the Marojejy National Park area to ThaiV and its descendants. The outgroup (black) is a Sangassou virus. Support level shows posterior probability. Nodes with a posterior probability >0.75 are represented with black squares while red squares represent nods with probabilities >0.95. Specimens from Mandena are in yellow, Manantenina in red, and Sarahandrano in blue .and all are from Rattus rattus. Accession numbers are listed in supplementary Table S5. Satellite images were produced using Google SNES/Airbus Maxar Technolgies Data SIO, NOAA, U.S. Navy, NGA, GEBCO and the north indicated.

### 3.1. Relationship between environmental factors and hantavirus prevalence

Infected *R. rattus* were present in all habitat types with the exception of semi-intact forests within the park where all 52 individuals tested negative. Across the other habitat types, prevalence differed significantly (p=0.001, Fisher’s exact test, Fig. 1). The prevalence was also significantly different by season (p=0.01, Fisher’s exact test) and study site (p<0.001, Fisher’s exact test) but did not differ by distance to village (p=0.595, Wilcoxon test). Hantavirus prevalence in a grid correlated positively with the number of *R. rattus* captured on that grid (p<0.001, ρ=0.49, Spearman’s rank correlation). Because the trap grid design was altered to include 21 additional traps starting in September 2019, we verified that no significant differences occurred in the average number of *R. rattus* captured per grid before and after this change (Wilcoxon test, p=0.866). Across trap grid installations (omitting trapping done in homes in the village), the average number of *R. rattus* captured per grid was 16.3 ± 11.2 (range 1 to 42).

The best-models subset for the environmental covariates (habitat type, village, season, and distance to village) contained a model with habitat type, season, and village (AICc weight 0.501); a model with habitat type and village (AICc weight 0.310); and the global model with all variables (AICc weight 0.189). The most important predictors were habitat type and village, which were present in all models (AICc weights 1.00). The season and distance to the village had respective weights of 0.69 and 0.19. The full model-averaged estimates are shown in Fig. 3 and Table S4. Post-hoc comparisons between habitat types (Fig. 3) revealed that the probability of infection was highest in flooded rice fields (0.1607) and agroforests (0.1277), and lowest in secondary forests (0.0403) and villages (0.0262). The prevalence between flood rice fields and village habitat types was significantly different (p=0.030). Semi-intact forests were excluded from this analysis because there were no positive animals. The probability of infection in Sarahandrano (0.1821) was significantly higher than in both Mandena (0.0501, p<0.001) and Manantenina (0.0444, p=0.033). By contrast, there was no significant difference in the probability of infection between the three seasons (p>0.05).

**Figure 3.**
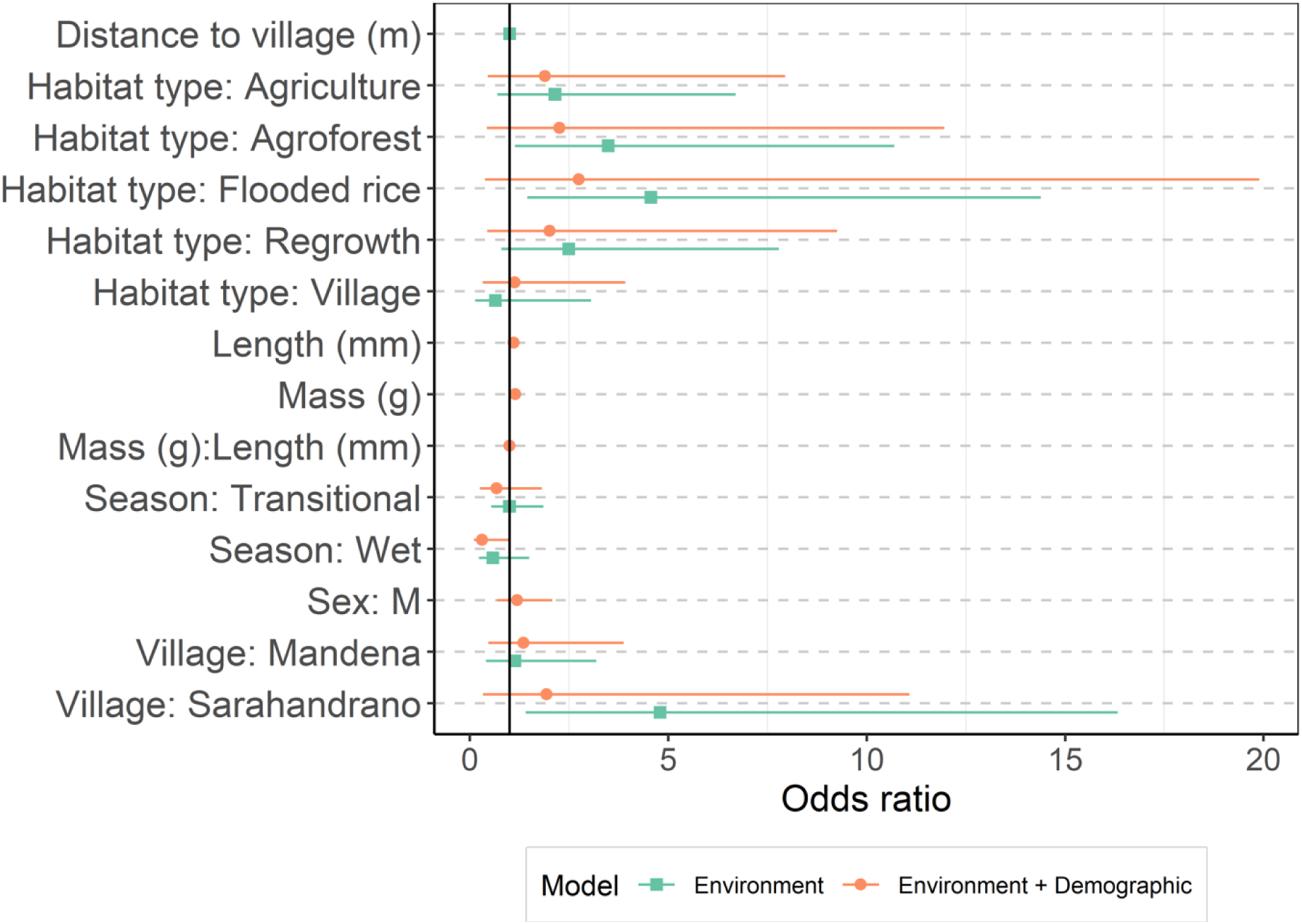
Estimated coefficients of the GLMMs. Full model-averaged coefficients for the GLMM of models containing environmental (green squares) and environmental as well as demographic (orange circles) predictors. The coefficients (shapes) are shown with 95% confidence intervals. Fig S3. Shows comparisons of all four models considered.

**Figure 4.**
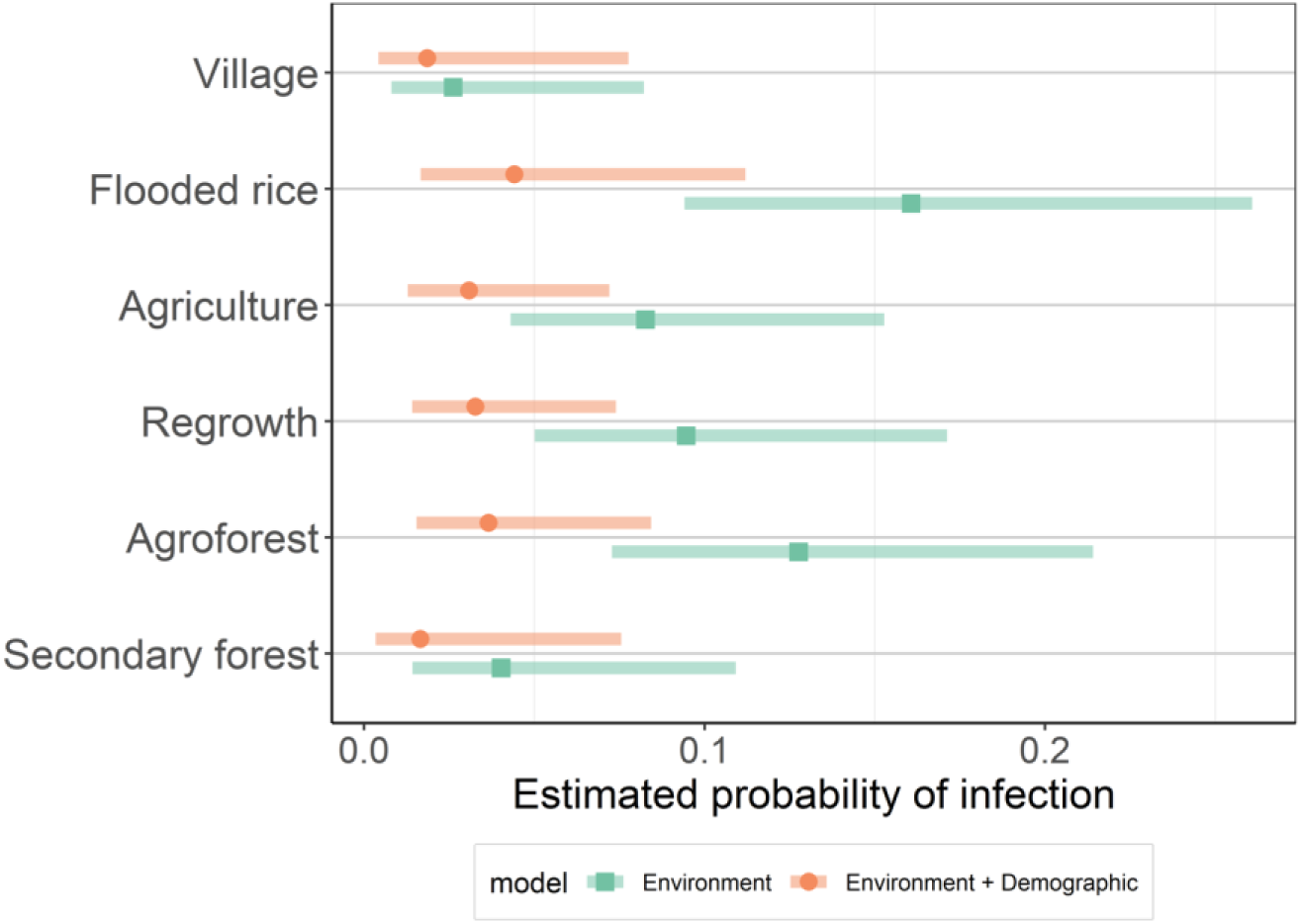
Predicted probability of hantavirus infection in R. rattus across habitat types. Estimated marginal means probability of hantavirus infection across the different habitat types after accounting for variability in prevalence due to other variables in the respective models, with a Tukey adjustment for multiple comparisons. The post hoc estimate made by the environment model (green square) and environment + demographic model (orange circle) and shaded areas of their corresponding 95% confidence intervals are displayed. Fig S4 shows comparisons of all 4 models considered and other categorical predictors.

### 3.2. Relationship between *Rattus rattus* demography and hantavirus prevalence

Hantavirus was not detected in any of the sub-adult *R. rattus* (N=137, Fig. 1). The mass of *R. rattus* in which we detected hantavirus was 50% heavier (mean = 125.0 ± 28.1g) than those testing negative (81.7 ± 41.3g; Welch’s two sample t-test; p<0.001; Fig. 1b). The infected *R. rattus* also had significantly greater head-body length (166.2 ± 22.8 mm vs 139.0 ± 34.0 mm; p <0.001; Welch’s two sample t-test), but not significantly different body condition scores (p=0.470, Welch’s two sample t-test). Prevalence in males (12.3%, 51/416) was significantly higher than in females (6.1%, 23/375; p=0.005, χ^2^=8.02).

The best model subset of the 12 possible models of individual-level predictors of infection included a model with the interaction between age and mass, and the lower order terms (AICc weight 0.435) and four other models with AICc weights <0.3 (Fig. S3 and Table S4). Mass and head-body length were the most important predictors and were present in all models (cumulative sum weight = 1) followed by their interaction (cumulative sum weight =0.97). Sex and body condition score were both in two models, with sum weights of 0.37 and 0.26 respectively.

### 3.3. Covariance between habitat type and demography

The morphometric measurements of trapped *R. rattus* covaried with the environmental predictors of infection, namely habitat types, seasons and villages. Body mass and head-body length were significantly different across all three variables (p<0.001, Kruskal-Wallis) while body condition was also significantly different across habitat types and seasons (p=0.047, Kruskal-Wallis), but not villages (p=0.64, Kruskal-Wallis). Age was significantly different across habitat types (p<0.001), seasons (p=0.005) and villages (p<0.001, Fisher’s exact test). By contrast, sex ratio was not significantly different across habitat types (p=0.56), seasons (p=0.90), or villages (p=0.01, Fisher’s exact test). Distance to village was not strongly correlated with mass or body condition and was not significantly different between ages or sexes (p=0.21 and 0.08; respectively, Wilcoxon test). Full-model averaged coefficients and the importance of predictors is provided in Table S4.

## 4. Individual and environmental predictors of infection

To address whether the observed differences in the probability of infection between habitat types were due to demographic variability, rat density, or a combination of these effects, three additional GLMM averaged results were considered. The best fit GLMM combined environmental (habitat types, seasons, and villages) and demographic (mass, head-body length, their interaction, and sex) effects. Full model averaged coefficients are shown in Fig. 2, Fig. S3, and Table S4. The most important predictors across the 16 models in the 95% cumulative sum weight subset were head-body length and mass (sum weights of 1), followed by their interaction term (14 models; sum weight = 0.98). The next important effects were season (10 models, sum weight = 0.90), habitat type (6 models, sum weight = 0.57), village (7 models, sum weight = 0.46) and sex (7 models, sum weight = 0.44). Post-hoc comparisons of the probability of infection between habitat types show that the effect of habitat type is minimized when demographic effects are included in the model (Fig. 3). The descriptions and estimated full-model averaged coefficients of the GLMMs containing (i) environmental effects along with the number of *R. rattus* captured and (ii) all predictors except body condition and distance to village can be found in Table S4, Fig S3, and Fig S4.

## Discussion

Our extensive investigation of hantavirus in small mammals from northeastern Madagascar only detected infections in *R. rattus* (9.5%) despite testing over 20 species of terrestrial mammals, composed of 1663 individuals (including 450 endemic specimens) and 227 native bats. The detected hantavirus is closely related to ANJZV/MayoV, which were previously described on Madagascar and Mayotte (Filippone et al., 2016; Rabemananjara et al., 2020; Reynes et al., 2014). Infection prevalence was highest in animals captured in agricultural land use areas. This effect weakened, however, when the variability in animal body size between land-use types was taken into account. The findings involving body size highlight the importance of demographic factors in reservoir populations for understanding prevalence and spillover risk, while also identifying potential mechanisms that relate land use change to zoonotic disease risk.

The absence of hantavirus detection in any of the endemic or native species may be due to extremely low prevalence or the presence of a distant viral lineage that could not be detected with our PCR scheme. We screened 269 of the 450 trapped endemic terrestrial mammals with three alternative semi-nested PCR schemes, which still led to negative results (Supplementary Table S1). ANJZV has been previously reported from the endemic rodent *Eliurus majori* (n tested = 15) in the Central Highlands of Madagascar with the same PCR scheme used herein (Reynes et al., 2014). Since hantaviruses are notorious for host switching (Guo et al., 2013), we can hypothesize that the absence of detectable virus in endemic animals mirrors a recent introduction to Madagascar, where adaptation to the endemic species is in an early stage. However, further investigations are needed to date the introduction of the virus.

The viral sequences were distinct but genetically nested within the clade containing ANJZV, MayoV, and ThaiV sequences. The viral lineage found in *R. rattus* from our samples in the Marojejy National Park area in northeast Madagascar show less than 92% identity with ANJZV from central eastern Madagascar, located 480 km southwest of the study site, and MayoV from Mayotte Island, located 530 km northwest of the study site. However, longer sequences are needed to robustly establish the genetic relationships of the virus occurring in the Marojejy area and the previously reported ANJZV and MayoV.

Prevalence of hantavirus infection in our study was similar to other studies of the virus on Madagascar (Raharinosy et al., 2018; Reynes et al., 2014). We found an infection prevalence of 9.5% in *R. rattus*, which is not significantly different from a previous study reporting an infection prevalence in *R. rattus* of 12.4% across the island (χ^2^=2.7, p=0.10) and 5.8% in Sambava (χ^2^=0.69, p=0.41), in close proximity to our study site (Raharinosy et al., 2018). Notably, both of these studies predicted higher prevalence in the more mesic regions of Madagascar (Raharinosy et al., 2018), such as where the present study took place, which we did not observe (Raharinosy et al., 2018). Our findings which did not demonstrate strong seasonal trends in hantavirus prevalence, as predicted by Raharinosy et al. (2018), align with the mild seasonal patterns of this region of Madagascar.

Our sampling schema across a matrix of land-use types allowed us to highlight determinants of infection at a finer scale than previous studies (Raharinosy et al., 2018; Reynes et al., 2014). We found that agricultural land-use types (agroforest, brushy regrowth, agriculture, and flooded rice fields) displayed higher prevalence than the most disturbed (village) and least disturbed (semi-intact and secondary forest) habitats. Of note, not a single *R. rattus* captured in the semi-intact forest (n=52) tested positive. The low prevalence in the village is in accordance with previous findings from Madagascar (Raharinosy et al., 2018). The overall effect of habitat type on infection probability appears to be due to differences in individual traits, particularly the size (mass and length) and sex of *R. rattus* captured in each habitat type. In the Manantenina study area, statistically significant differences in body size of *R. rattus*, based on cranio-dental measurements, have been identified between native forest and anthropogenic habitats (Ranaivoson et al., 2022). The niche breadth of animals living in natural forest was greater than in anthropogenic habitats (Dammhahn et al., 2017), presumably indicating a more stable diet during periods of seasonal variation and driving higher life expectancy. Hence, the longer-lived animals have a greater chance to come in contact with the virus. Alternatively, we cannot exclude that older males tend to leave the villages or are more at risk to be killed by domestic animals and people, hence contributing to lower overall infection prevalence.

Based on these findings, we propose that differences in hantavirus prevalence in *R. rattus* across our habitat gradient is due to a succession of synergetic biotic and abiotic factors, including resource availability, animal density, life expectancy, and level of habitat disturbance. Together these factors impacted population demography, which appears to drive infection prevalence in this system. Disturbances that alter the population demography to favor larger-bodied and presumably older individuals may lead to increases in prevalence and thus human exposure risk. Based on the results that prevalence was highest in agricultural fields, we also expect that human exposure risk is highest when conducting farm-based activities that aerosolize excreta or bring people into contact with *R. rattus*.

## Conclusions

*Rattus rattus* is a remarkably adaptable species, and appears to be the principal reservoir of hantavirus in the area surrounding Marojejy National Park in northeastern Madagascar. The positiuve individuals from our study formed a subclade of MayoV and Anjozorobe strains of Thailand hantaviruses, both of which have previously been described on Madagascar. Infection prevalence in *R. rattus* varied across the land-use matrix and was higher in agricultural areas than in forests and villages. Highly disturbed habitats had higher abundances of *R. rattus*. However, the larger body size of *R. rattus* living in agricultural land-use types compared forests and villages likely explained the increased viral prevalence. Differences in body size likely indicate a longer lifespan and increased odds of viral exposure. As invasive rat populations and interactions between endemic and introduced species continue to grow due to the ongoing conversion of forest into agricultural land, we expect hantavirus exposure risk to humans to increase, particularly when these changes positively alter *R. rattus* demography.

## Acknowledgement

This research was funded by the joint NIH-NSF-NIFA Ecology and Evolution of Infectious Disease award R01-TW011493, the Duke Global Health Institute, the Bass Connections Program at Duke University, and a Duke University Provost’s Collaboratory grant. In addition, the collaboration between Duke University and the Universite de La Réunion was supported by the Thomas Jefferson Fellowship from the FACE Foundation; http://facefoundation.org/thomas-jefferson-fund/. We are grateful to the Mention Zoologie et Biodiversite Animale, Université d’Antananarivo; Madagascar National Parks; and the Direction des Aires Protégées, des Ressources Naturelles Renouvelables et des Ecosystèmes for administrative aid and issuing research permits. We thank the PIMIT laboratory and all the technical assistance given by Magalie Turpin. The Duke Lemur Center’s SAVA Conservation Initiative provided logistical and other support in the SAVA Region. We also thank those who have hosted us in their communities over the years of trapping that occurred, including the many field assistants from the villages where research took place.

## Authors contributions

The project was conceptualized by PT, CLN, JPH, SMG, and VS. Funding Acquisition was done by PT, CLN, SMG and JPH. Field investigations were carried out by TMR and JPH. Laboratory investigators were JD and PT. Data Curation was done by KMK, JPH, and TMR. Formal analysis and writing of the original draft was done by JD and KMK. All authors reviewed and edited the work.

## References

Avšič-Županc, T., Saksida, A., Korva, M., 2019. hantavirus infections. Clin. Microbiol. Infect. 21, e6–e16.

Bartoń K (2023). MuMIn: Multi-Model Inference. R package version 1.47.5, <https://CRAN.R-project.org/package=MuMIn>.

Brooks, M.E., Kristensen, K., Van Benthem, K.J., Magnusson, A., Berg, C.W., Nielsen, A., Skaug, H.J., Machler, M., Bolker, B.M., 2017. glmmTMB balances speed and flexibility among packages for zero-inflated generalized linear mixed modeling. R J. 9, 378–400.

Cascio, A., Bosilkovski, M., Rodriguez-Morales, A. J., & Pappas, G. (2011). The socio-ecology of zoonotic infections. Clinical microbiology and infection, 17(3), 336–342.

Clement, J., LeDuc, J.W., Lloyd, G., Reynes, J.-M., McElhinney, L., Van Ranst, M., Lee, H.-W., 2019. Wild rats, laboratory rats, pet rats: Global Seoul hantavirus disease revisited. Viruses 11, 652.

Dammhahn, M., Randriamoria, T.M. & Goodman, S.M. 2017. Broad and flexible stable isotope niches in invasive non-native Rattus spp. in anthropogenic and natural habitats of central eastern Madagascar. BMC Ecol.,17 (1), 6. 10.1186/s12898-017-0125-0.

Fialho, M.Y.G., Cerboncini, R.A.S., Passamani, M., 2019. Linear forest patches and the conservation of small mammals in human-altered landscapes. Mamm. Biol. 96, 87–92. 10.1016/j.mambio.2018.11.002

Filippone, C., Castel, G., Murri, S., Beaulieux, F., Ermonval, M., Jallet, C., Wise, E.L., Ellis, R.J., Marston, D.A., McElhinney, L.M., Fooks, A.R., Desvars, A., Halos, L., Vourc’h, G., Marianneau, P., Tordo, N., 2016. Discovery of hantavirus circulating among Rattus rattus in French Mayotte Island, Indian Ocean. J. Gen. Virol. 10.1099/jgv.0.000440

Ganzhorn, J.U., 2003. Effects of introduced Rattus rattus on endemic small mammals in dry deciduous forest fragments of western Madagascar. Anim. Conserv. 6, 147–157. 10.1017/S1367943003003196

Goodin, D.G., Koch, D.E., Owen, R.D., Chu, Y.-K., Hutchinson, J.M.S., Jonsson, C.B., 2006. Land cover associated with hantavirus presence in Paraguay: Land cover associated with hantavirus presence. Glob. Ecol. Biogeogr. 15, 519–527. 10.1111/j.1466-822X.2006.00244.x

Goodman, S.M., 2022. The new natural history of Madagascar. Princeton University Press.

Guo, W.-P., Lin, X.-D., Wang, W., Tian, J.-H., Cong, M.-L., Zhang, H.-L., Wang, M.-R., Zhou, R.-H., Wang, J.-B., Li, M.-H., Xu, J., Holmes, E.C., Zhang, Y.-Z., 2013. Phylogeny and Origins of hantaviruses Harbored by Bats, Insectivores, and Rodents. PLOS Pathog. 9, e1003159. 10.1371/journal.ppat.1003159

Hartig, F., 2022. DHARMa: Residual Diagnostics for Hierarchical (Multi-Level / Mixed) Regression Models. R package version 0.4.6.

Jonsson, C.B., Figueiredo, L.T.M., Vapalahti, O., 2010. A global perspective on hantavirus ecology, epidemiology, and disease. Clin. Microbiol. Rev. 23, 412–441.

Kang, H.J., Bennett, S.N., Dizney, L., Sumibcay, L., Arai, S., Ruedas, L.A., Song, J.-W., Yanagihara, R., 2009. Host switch during evolution of a genetically distinct hantavirus in the American shrew mole (Neurotrichus gibbsii). Virology 388, 8–14. 10.1016/j.virol.2009.03.019

Kikuchi, F., Senoo, K., Arai, S., Tsuchiya, K., Sơn, N.T., Motokawa, M., Ranorosoa, M.C., Bawm, S., Lin, K.S., Suzuki, H., Unno, A., Nakata, K., Harada, M., Tanaka-Taya, K., Morikawa, S., Suzuki, M., Mizutani, T., Yanagihara, R., 2021. Rodent-Borne Orthohantaviruses in Vietnam, Madagascar and Japan. Viruses 13, 1343. 10.3390/v13071343

Klempa, B., 2009. hantaviruses and climate change. Clin. Microbiol. Infect. 15, 518–523.

Klempa, B., Fichet-Calvet, E., Lecompte, E., Auste, B., Aniskin, V., Meisel, H., Denys, C., Koivogui, L., ter Meulen, J., Krüger, D.H., 2006. hantavirus in African Wood Mouse, Guinea. Emerg. Infect. Dis. 12, 838–840. 10.3201/eid1205.051487

Lenth R (2023). emmeans: Estimated Marginal Means, aka Least-Squares Means. R package version 1.8.6.

Nsoesie, E.O., Mekaru, S.R., Ramakrishnan, N., Marathe, M.V., Brownstein, J.S., 2014. Modeling to Predict Cases of hantavirus Pulmonary Syndrome in Chile. PLoS Negl. Trop. Dis. 8, e2779. 10.1371/journal.pntd.0002779

Pardini, R., Bueno, A. de A., Gardner, T.A., Prado, P.I., Metzger, J.P., 2010. Beyond the Fragmentation Threshold Hypothesis: Regime Shifts in Biodiversity Across Fragmented Landscapes. PLoS ONE 5, e13666. 10.1371/journal.pone.0013666

Prist, P.R., D Andrea, P.S., Metzger, J.P., 2017. Landscape, Climate and hantavirus Cardiopulmonary Syndrome Outbreaks. EcoHealth 14, 614–629. 10.1007/s10393-017-1255-8

R Core Team, R., 2023. R: A language and environment for statistical computing. R Foundation for Statistical Computing, Vienna, Austria <https://www.R-project.org/>

Rabemananjara, H.A., Raharinosy, V., Razafimahefa, R.M., Ravalohery, J.P., Rafisandratantsoa, J.T., Andriamandimby, S.F., Rajerison, M., Rahelinirina, S., Harimanana, A., Irinantenaina, J., Olive, M.-M., Rogier, C., Tordo, N., Ulrich, R.G., Reynes, J.-M., Petres, S., Heraud, J.-M., Telfer, S., Filippone, C., 2020. Human Exposure to hantaviruses Associated with Rodents of the Murinae Subfamily, Madagascar. Emerg. Infect. Dis. 26, 587. 10.3201/eid2603.190320

Raharinosy, V., Olive, M.-M., Andriamiarimanana, F.M., Andriamandimby, S.F., Ravalohery, J.-P., Andriamamonjy, S., Filippone, C., Rakoto, D.A.D., Telfer, S., Heraud, J.-M., 2018. Geographical distribution and relative risk of Anjozorobe virus (Thailand orthohantavirus) infection in black rats (Rattus rattus) in Madagascar. Virol. J. 15, 83. 10.1186/s12985-018-0992-9

Ranaivoson, T.N., James P. Herrera, Voahangy Soarimalala, Toky M. Randriamoria, and Steven M. Goodman. 2022. La variation morphologique de Rattus rattus Linnaeus, 1758 (Rodentia : Muridae) dans les habitats forestiers et anthropisés du bassin-versant nord-est de Madagascar. Bulletin de la Société Zoologique de France, 147 (3): 129–141.

Reynes, J.-M., Razafindralambo, N.K., Lacoste, V., Olive, M.-M., Barivelo, T.A., Soarimalala, V., Heraud, J.-M., Lavergne, A., 2014. Anjozorobe hantavirus, a new genetic variant of Thailand virus detected in rodents from Madagascar. Vector-Borne Zoonotic Dis. 14, 212–219.

Schmaljohn, C., Hjelle, B., 1997. hantaviruses: a global disease problem. Emerg. Infect. Dis. 3, 95–104. 10.3201/eid0302.970202

Scobie, K., Rahelinirina, S., Soarimalala, V., Andriamiarimanana, F.M., Rahaingosoamamitiana, C., Randriamoria, T., Rahajandraibe, S., Lambin, X., Rajerison, M., ℡fer, S., n.d. Reproductive ecology of the black rat (Rattus rattus) in Madagascar: the influence of density-dependent and -independent effects. Integr. Zool. n/a. 10.1111/1749-4877.12750

Sumibcay, L., Kadjo, B., Gu, S.H., Kang, H.J., Lim, B.K., Cook, J.A., Song, J.-W., Yanagihara, R., 2012. Divergent lineage of a novel hantavirus in the banana pipistrelle (Neoromicia nanus) in Côte d’Ivoire. Virol. J. 9, 34. 10.1186/1743-422X-9-34

Suzán, G., Marcé, E., Giermakowski, J.T., Armién, B., Pascale, J., Mills, J., Ceballos, G., Gómez, A., Aguirre, A.A., Salazar-Bravo, J., Armién, A., Parmenter, R., Yates, T., 2008. The Effect of Habitat Fragmentation and Species Diversity Loss on hantavirus Prevalence in Panama. Ann. N. Y. Acad. Sci. 1149, 80–83. 10.1196/annals.1428.063

Umetsu, F., Pardini, R., 2007. Small mammals in a mosaic of forest remnants and anthropogenic habitats—evaluating matrix quality in an Atlantic forest landscape. Landsc. Ecol. 22, 517–530. 10.1007/s10980-006-9041-y

Vapalahti, O., Lundkvist, Å., Fedorov, V., Conroy, C.J., Hirvonen, S., Plyusnina, A., Nemirov, K., Fredga, K., Cook, J.A., Niemimaa, J., Kaikusalo, A., Henttonen, H., Vaheri, A., Plyusnin, A., 1999. Isolation and Characterization of a hantavirus from Lemmus sibiricus : Evidence for Host Switch during hantavirus Evolution. J. Virol. 73, 5586–5592. 10.1128/JVI.73.7.5586-5592.1999

Watson, D.C., Sargianou, M., Papa, A., Chra, P., Starakis, I., Panos, G., 2014. Epidemiology of hantavirus infections in humans: a comprehensive, global overview. Crit. Rev. Microbiol. 40, 261–272. 10.3109/1040841X.2013.783555

Weiss, S., Witkowski, P.T., Auste, B., Nowak, K., Weber, N., Fahr, J., Mombouli, J.-V., Wolfe, N.D., Drexler, J.F., Drosten, C., Klempa, B., Leendertz, F.H., Kruger, D.H., 2012. hantavirus in Bat, Sierra Leone. Emerg. Infect. Dis. 18, 159–161. 10.3201/eid1801.111026

